# Microfluidic Electric Egg-Laying Assay and Application to *In-vivo* Toxicity Screening of Microplastics using *C. elegans*

**DOI:** 10.1101/2021.01.16.426931

**Authors:** Khaled Youssef, Daphne Archonta, Terrance J. Kubiseseki, Anurag Tandon, Pouya Rezai

**Author notes:** Corresponding Author: BRG 433B, 4700 Keele St, Toronto, ON, M3J 1P3, Canada.

## Abstract

Environmental pollutants like microplastics are posing health concerns on aquatic animals and the ecosystem. Microplastic toxicity studies using *C. elegans* as a model are evolving but methodologically hindered from obtaining statistically strong data sets, detecting toxicity effects based on microplastics uptake, and correlating physiological and behavioural effects at an individual-worm level. In this paper, we report a novel microfluidic electric egg-laying assay for phenotypical assessment of multiple worms in parallel. The effects of glucose and polystyrene microplastics at various concentrations on the worms’ electric egg-laying, length, diameter, and length contraction during exposure to electric signal were studied. The device contained eight parallel worm-dwelling microchannels called electric traps, with equivalent electrical fields, in which the worms were electrically stimulated for egg deposition and fluorescently imaged for assessment of neuronal and microplastic uptake expression. A new bidirectional stimulation technique was developed, and the device design was optimized to achieve a testing efficiency of 91.25%. Exposure of worms to 100mM glucose resulted in a significant reduction in their egg-laying and size. The effects of 1μm polystyrene microparticles at concentrations of 100 and 1000 mg/L on the electric egg-laying behaviour, size, and neurodegeneration of N2 and NW1229 (expressing GFP pan-neuronally) worms were also studied. Of the two concentrations, 1000 mg/L caused severe egg-laying deficiency and growth retardation as well as neurodegeneration. Additionally, using single-worm level phenotyping, we noticed intra-population variability in microplastics uptake and correlation with the above physiological and behavioural phenotypes, which was hidden in the population-averaged results. Taken together, these results suggest the appropriateness of our microfluidic assay for toxicological studies and for assessing the phenotypical heterogeneity in response to microplastics.

## INTRODUCTION

The discharge of environmental toxicants such as plastics, pesticides, carcinogens, antimicrobial products, and neurotoxins to the environment is escalating, which poses a significant burden on society, and the ecosystem’s health and safety.^1-3^ Some of these toxicants have been linked to neurodegeneration effects and reproductive complications.^4,5^ A total of 359 million metric tons of plastics were manufactured from 1950 to 2018^6^, leading to a severe accumulation of plastic waste in the environment. Plastic debris of less than 5 mm in size, termed microplastics^1^, has become a major concern because it can travel over long distances in the air and aquatic environments while being easily ingested by organisms and animals.^3,5,7,8^

The effects of microplastics and their characteristics of shape, size, and concentration on marine life and the ecosystem have been studied and shown to result in various dose-dependent toxicities, including neural, behavioural, and reproductive toxicity.^9-12^ The use of mammalian animal models limits the sample size in these studies and weakens their statistical strength. Such assays are also laborious, expensive, and time-consuming.^13,14^ Rapid and high-throughput *in-vivo* screening assays for microplastic toxicity studies are urgently needed. In this front, simple model organisms such as *Caenorhabditis elegans, Drosophila melanogaster*, and *Danio rerio* have been used for toxicological studies to rapidly examine the effects of different pollutants at throughputs higher than animal-based assays.^15-18^

*C. elegans* has been used as a model organism for studying microplastics toxicity, offering versatile experimental advantages including small size, short life cycle, ease of maintenance, biological simplicity, and body transparency, leading to its suitability for genetic modification and high-throughput cell-to-behaviour toxicity screening.^19-21^ Ingestion and intestinal accumulation of microplastics smaller than 5μm have been shown in *C. elegans*.^10,22,23^ Microplastics negatively affected *C. elegans* phenotypical behaviours such as locomotion and body bend frequency as well as its growth and reproduction.^9-12,22-24^

Lei et al.^11^ studied the effects of exposing *C. elegans* to 1 mg/L of 0.1-5μm polystyrene microparticles, using locomotory behaviours, growth, and lifespan as toxicity indicators. Their investigations demonstrated that 1μm particles significantly deteriorated the survival and growth rate and caused oxidative damage to cholinergic and GABAnergic neurons, which was ameliorated by natural antioxidants such as curcumin. The same group investigated the toxic effects of different microplastics, including polyamides, polyethylene, polypropylene, polyvinyl chloride, and polystyrene at various concentrations of 0.5-10 mg/m^2^.^12^ All microplastics showed similar toxic effects on the survival rate of *C. elegans*, indicating no apparent dose-or material-dependent relationship. However, different sizes of 0.1-5μm microparticles showed a size-dependent lethality with a significant decrease in the reproduction rate and brood size at 1μm microparticles. Yu et al.^22^ investigated the toxicity mechanisms of 1 μm polystyrene microplastics on *C. elegans* at different concentrations of 0-100 μg/L. Exposure to concentrations higher than 10μg/L induced a significant reduction in worms’ head thrash, body bend, body length, and brood size. The toxicity was attributed to the increased reactive oxygen species expression and intestinal damage. The above-mentioned studies showed the effects of microplastics at relatively low concentrations. Therefore, Kim et al.^25^ exposed *C. elegans* to 42 and 530 nm polystyrene particles at higher concentrations up to 100 mg/L and showed a significant reduction in the brood size. The effect of microplastics on other behaviours of *C. elegans*, such as reproduction, remains largely unknown. Another gap is the heterogeneity of microplastics uptake by the worms and a lack of understanding of correlating phenotypic toxicities, which requires the use of single-worm assays.

*C. elegans* reproduction is a rhythmic activity that has been established as a robust readout for investigating the toxic effects of different materials.^22,26^ The egg-laying rate is measured by performing progeny counting over several hours. The entire process takes up to 8 hours and requires worm picking expertise without affecting its health, followed by counting the eggs. This conventional technique is cumbersome and limited in throughput to a few worms per hour. To address this limitation, we recently reported a novel technique to stimulate egg-laying of adult *C. elegans* on-demand using electric pulses in a microchannel (termed electric egg-laying).^27,28^ The throughput of our device was limited to 5 worms/hr, hindering the feasibility of using it in toxicological studies on 100s of worms/assay.

In this paper, we report a new microfluidic device for electric egg-laying analysis and on-chip fluorescent imaging of multiple worms in parallel and apply it for the first time to toxicity screening of microplastics. The established effect of glucose on natural reproduction^29^ was used as a proof-of-principle experiment to test the suitability of our assay. Then, we demonstrated the novel application of our method for microplastic toxicity studies, showing the adverse effects of microplastics on the electric egg-laying behaviour of *C. elegans* and its correlation with microplastic accumulation in the worms. We achieved an assay time of 10 min and a throughput of up to 40 worms/hr, which was mainly limited by our microscopic field of view. Our method is significantly faster than conventional egg-laying assays with 4 hr assay time and 5 worms/hr throughput. Moreover, we showed another interesting advantage of our device for investigating the effect of microplastics uptake in correlation with multiple physiological and behavioural phenotypes of *C. elegans*, all at a single-worm resolution, which is not readily achievable by conventional methods.

## MATERIALS AND METHODS

### *C. elegans* culturing

Wild type *C. elegans* strain was obtained from the *Caenorhabditis* Genetics Center (University of Minnesota, USA) and maintained at approximately 22°C on freshly prepared standard nematode growth media (NGM) plates with *Escherichia coli (E. coli)* strain *OP50* as a food source.^30^ All assays were performed with well-fed gravid hermaphrodite worms (day one post young adult stage (~64hrs)) obtained using the conventional alkaline hypochlorite (bleach) treatment.^31^ Briefly, one week prior to the experiments, a chunked NGM plate was prepared and left for three days until a majority of the worms reached the gravid adult stage. All worms were washed off the plate in a 15 mL Eppendorf tube and exposed for 10 min to a solution of 3.875 mL double-distilled water, 125 μL NaOH, and 1 mL commercial bleach. The eggs were then collected by washing off the bleach solution using double-distilled water and centrifuging at 1500 rpm. The collected eggs were allowed to hatch into L1 larvae overnight in 1 mL of M9 buffer (3 g KH_2_PO_4_, 6 g Na_2_HPO_4_, 5 g NaCl, and 1 mL 1 M MgSO_4_ in 1 L distilled water) using a RotoFlex™ tube rotator (RK-04397-40, Cole-Parmer, Canada). L1 larvae were collected by centrifuging and seeded on top of NGM plates, containing glucose or microplastics when needed.

### *C. elegans* exposure to glucose and microplastics

Glucose was used as an established toxicant to natural egg-laying^29^ to test the performance of our multi-worm device before using microplastics. NGM plates were prepared 5 days before the experiment, either with or without 100mM of glucose, and left for three days before bacterial seeding.^29^ Luria Broth (LB) media (10 g Bacto-tryptone, 5 g Bacto-yeast, and 5 g NaCl in 1 L distilled water) was inoculated with a single colony of OP50 and cultured overnight at 37°C in a thermal shaker-incubator.^30^ Then, the NGM plates with or without glucose were seeded with 100 μL of the freshly prepared bacteria culture and left for two days before being used. L1 larvae were seeded on top of NGM plates with or without glucose for ~64 *hr* at 22°C until they become gravid adults and ready for our experiments.

Some studies^11,12^ have shown that the toxicity of microparticles *on C. elegans* is size-dependent rather than material-or dose-dependent, and 1μm particles have been shown to be an effective size. Our microplastic exposure protocol was based on the method introduced by Schöpfer et al.^24^. One percent (w/v) stock solutions of 1-1.4μm high-intensity polystyrene Nile red fluorescent microparticles were procured from Spherotech Inc (FH-1056-2, Lake Forest, USA). Microplastic feed suspensions were prepared at concentrations of 100 and 1000 mg/L in a mixture of M9 buffer and *E. coli OP50*. Aliquots of 100 μL of the prepared solutions were seeded on top of bacterial lawns on NGM plates. Then, the microplastic-seeded plates were left to dry for two days before being used to expose the worms. The microplastics distribution was confirmed to be uniform on top of the bacterial lawn with a fluorescent microscope (Leica MZ10F fluorescence microscope, Leica, Wetzlar, Germany). For control experiments, plates were seeded only with 100 μL of the *E. coli* in M9 buffer. The prepared plates were then used for *C. elegans* growth from the L1 stage to the gravid adult stage (~64 hr) at 22°C for our experiments.

### Off-chip egg-laying assay

The off-chip egg-laying assay was conducted for the glucose experiments as a validation step to prove that glucose was affecting the worms.^29^ On the day of the experiment, 25 worms per group (control and glucose exposed) were randomly picked and seeded on new NGM plates with and without glucose, respectively. After three hours, the worms were picked off the plates, and the number of laid eggs was counted using an inverted microscope (BIM-500FLD, Bio-imager Inc., Canada) equipped with a camera (SN 14120187, Point Grey Research Inc. Canada). Counting the eggs on a plate took 60 min on average. The obtained number of eggs was compared to the control egg-count.

### Microfluidic chip design and fabrication

The microfluidic device shown in Figure 1A was developed to enhance the throughput of our recently reported electric egg-laying method^27,28^, then used for glucose and microplastic toxicity studies.

**Figure 1:**
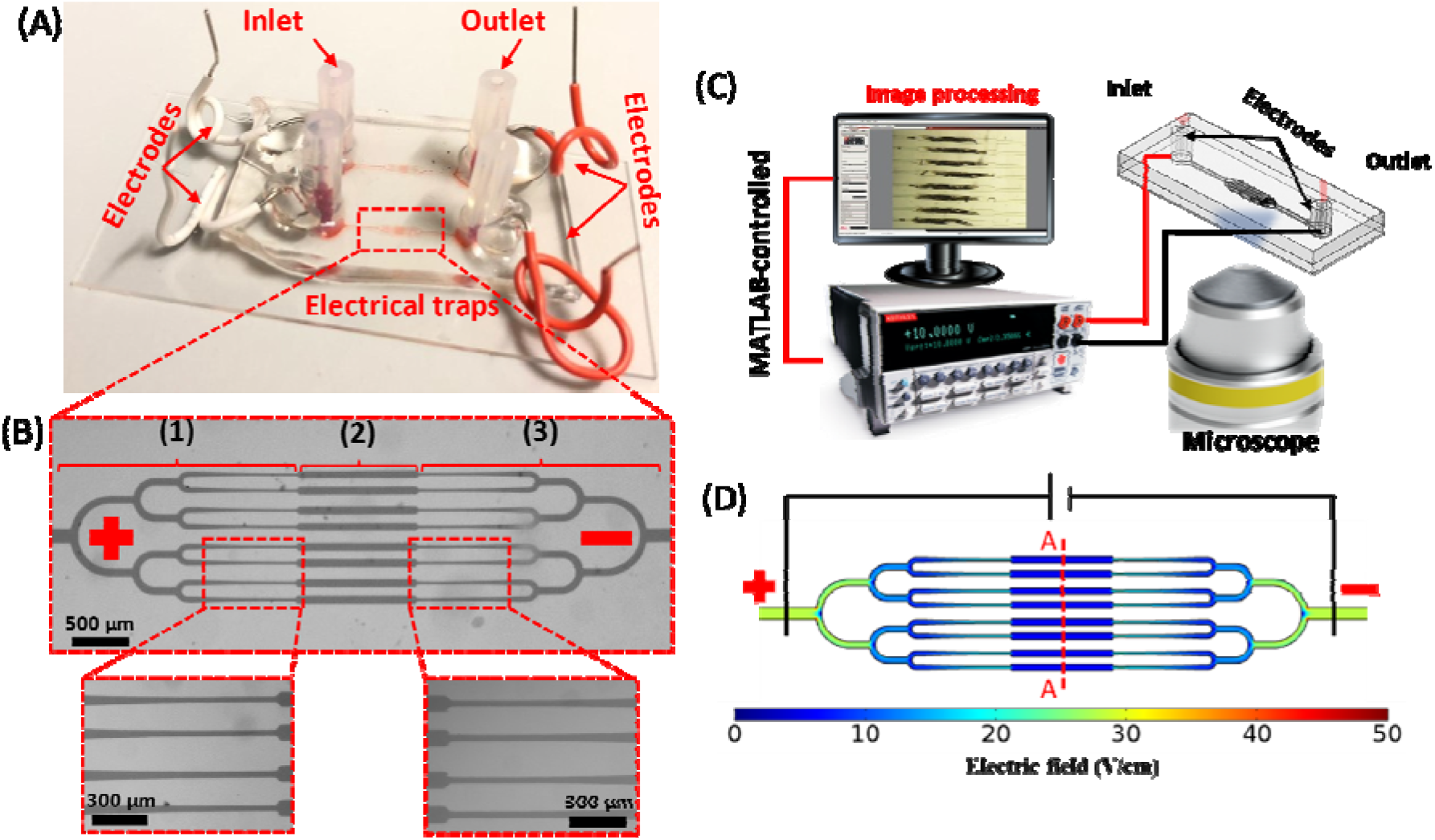
Microfluidic device and experimental setup for studying the electric egg-laying of 8 worms in parallel. (A) An image of the actual chip, housing two devices in parallel, each consisting of one inlet and one outlet interconnected with three-channel sections shown in (B): (1) Tree-like worm loading and distribution channels with end-tapered channels (left inset), (2) 8 parallel electrical traps for worm housing and imaging during the experiment, and (3) worm unloading channels with tapered connections to the electric traps (right inset). (C) Schematic of the experimental setup composing of the microfluidic device mounted on an inverted microscope and connected to a sourcemeter controlled by MATLAB. (D) COMSOL Multiphysics simulation of the electric field distribution throughout the chip to obtain a constant electric field of 6 V/cm in the electric traps using V= 34 Vat the inlet-outlet electrodes.

The 75 μm thick polydimethylsiloxane (PDMS) channel network in Figure 1B consisted of three-channel segments, i.e., (1) branching channels for worm loading and distribution with tapered endings (left inset of Figure 1B); (2) eight parallel 85 μm-wide and 1.3 mm long electric traps for worms’ egg-laying and imaging; and (3) branching channels for worm unloading and collection. The tapered ends of the loading channels started from a width of 60 μm and narrowed down to different widths of 30, 35, or 40 μm in three different device designs. The electrical traps were located at the mid-section of the device to provide a symmetrical electric field (EF) distribution delivered through the application of a DC voltage between the two metal wires in the inlet and outlet reservoirs.

The tree-like loading and unloading channels were inspired by Hulme et al.^32^ to ensure smooth loading and equal distribution of the worms across the 8 channels and maintain equal EF distribution. The loading and EF stimulation techniques benefit from the concept of maintaining constant channel dimensions at each bifurcation to preserve the same pressure and voltage drop up to the electrical traps. The pressure and voltage drops, as well as the hydrodynamic and electrical resistances, can be estimated using Hagen-Poiseuille’s (Eq. 1) and Ohm’s law (Eq. 2), respectively.

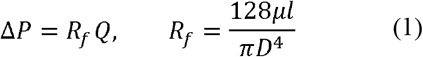

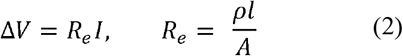

where *Q* is the flow rate, *R_f_* is the fluid flow resistance, *D* is the channel hydraulic diameter, *l* is the channel length, *μ* is the fluid dynamic viscosity, *A* is the cross-sectional area, *R_e_* is the electrical resistance, *I* is the electric current, and *ρ* is the electric resistivity.

Standard photo^33^- and soft-lithography^34^ techniques were used to fabricate the microfluidic device. At first, a SU-8 master mold was fabricated on a 4 in diameter and 500 μm thick silicon wafer (Wafer World Inc., USA) by UV (365nm with 11.1mW/cm^2^) exposure (UV-KUB 2, KLOE, France) of a 75 μm thick layer of SU-8 2075 photoresist (MicroChem Corporation, USA) for 20s. Then, to fabricate the negative replica of the master mold, a 10:1 mixture of PDMS elastomer and curing agent (Dow Corning, USA) was degasified, poured over the SU-8 mold, after placing two Masterflex tubes (L/S 14 size, Gelsenkirchen, Germany) over the inlet and outlet reservoirs, and cured for 2 hours at 80°C. An oxygen plasma machine (PDC-001-HP Harrick Plasma, USA) was used to bond the cured PDMS layer to a glass substrate at 870 mTorr pressure and 30 W power for 30s. Once the device was bonded, the two electrodes were punched through the inlet and outlet tubing for voltage application. Finally, to prevent leakage, the electrode punched areas were sealed with PDMS prepolymer and left to cure at 150°C for 5 minutes.

### Microfluidic egg-laying assay

Figure 1C depicts the experimental setup used to investigate the electric egg-laying of multiple worms in parallel and simultaneously image them at a single animal resolution. The experimental setup consisted of (1) our microfluidic device installed on a Leica inverted microscope (DMIL LED Inverted Routine Fluorescence Microscope, Leica, Germany) and imaged using a colour camera (MC170 HD, Leica, Germany), and (2) a direct current sourcemeter (Model 2410, Keithley Instruments Inc., USA) connected to the two electrodes at the inlet and outlet of the microfluidic device. The camera was used for recording movie clips and fluorescent images which were analysed using a custom-made MATLAB code. The code was also used to control the sourcemeter and apply the desired EF pulses while changing the EF polarity when needed. According to our single channel-based egg-laying experiments, an EF of 6 V/cm was needed to maximize egg-laying in *C. elegans*^22,35^ The required voltage to obtain this EF in the new microfluidic device was estimated to be 34 V using a simple COMSOL Multiphysics simulation (Figure 1D), as described in the Supplementary file Section S1.

To make our device accessible by the end-users, we eliminated the use of any fluidic control units such as syringe pumps, and the worms were manually manipulated using syringes. Briefly, a syringe filled with M9 buffer was connected to the inlet and used to fill in the device. Eight one day old worms were picked up manually from a NGM plate and loaded into the inlet. The worms were pressure-pulsed towards the electrical traps (Figure 2A) (Supplementary Video S1), and the loading efficiency (Eq. 3) was quantified and compared between the three devices with various loading tapered channel sizes.

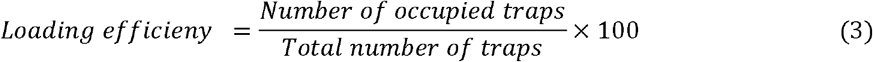

**Figure 2:**
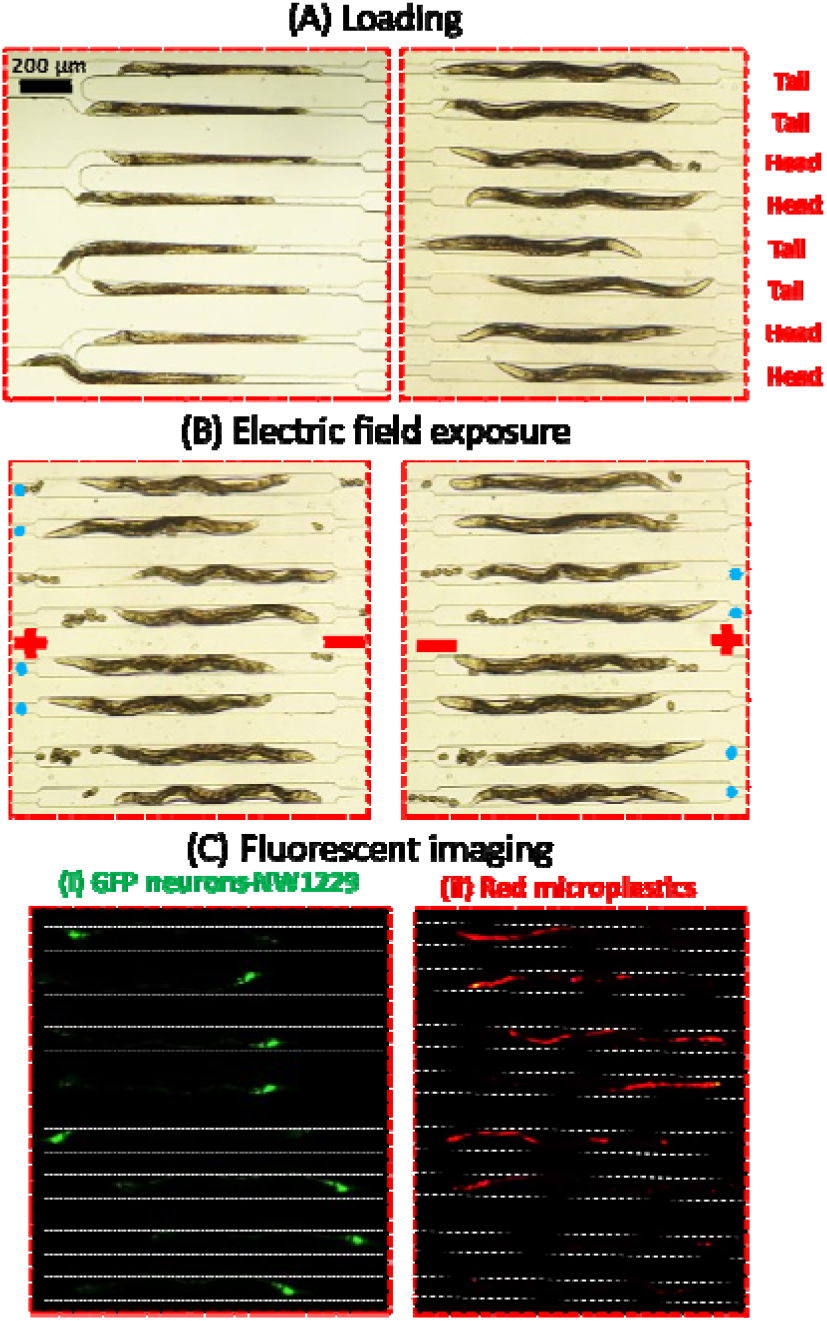
Capturing and investigating the electric egg-laying behaviour and microplastic accumulation in 8 worms in parallel. (A) Loading procedure through the branching channels resulting in a 50% tail vs. head orientation in the electric traps. (B) Egg-laying response of worms when the anode was at their head or tail. The eggs observed in the traps were deposited mostly when the worms were exposed to anode at their head, as donated by a blue circle beside them. (C) fluorescent images at the traps showing (i) GFP neurons of NW1229 worms and (ii) red fluorescent microplastics ingested by the same worms in (i). Channel walls are highlighted with white dashed lines for better observation.

Prior to EF stimulation, a 60 s acclimation period was used during which the flow was stopped by maintaining the inlet and outlet tubes at the same height level. Our previous experiments showed that the anode-facing worms could deposit significantly more eggs^27,35^, highlighting that the worm loading orientation with respect to the EF direction could affect our results. To maintain an equal exposure condition among randomly oriented worms in our device (as shown in Figure 2A), we adopted a new EF stimulation technique in which a series of 5s anodal and 5s cathodal pulses, separated by 25s acclimation periods, was applied for 10 min (Figure 2B). This ensured that each worm was stimulated with 10 anodal and 10 cathodal pulses during an experiment. Next, fluorescent imaging was conducted at 5x magnification for determining the accumulation of microplastics in each worm that was tested in the device (Figure 2C) (Supplementary Video S1). Then, the worms were flushed out of the chip for the next round of worms to be loaded.

### Data acquisition and analysis

The results, including the electric egg-count, the worms’ length, diameter, and length reduction during EF exposure, as well as mean fluorescent intensity expression of neurons (GFP) and microplastics (RFP), were extracted from the recorded videos using our custom-written MATLAB code.^27^ A step was added at the beginning of the code to select the 8 regions of interest around the worms in the electric traps. These regions were cropped, and the worms were individually analyzed, as shown in Figure S1.

We used two techniques for presenting our population-based data, i.e., bar plots with the mean ± standard error of the mean (SEM) and box plots with medians, 25% and 75% percentiles, and maximum and minimum data points. The population-based results were reported only for the worms responding to EF. The statistical significance between any two groups was determined using the Mann-Whitney test, while the following representation was used for identifying the significance level, i.e., * for p-value<0.05, ** for p-value<0.01, *** for p-value<0.001, and **** for p-value<0.0001.

Hierarchical Cluster Analysis (HCA) was performed to understand the toxicity effects of microplastics at single worm resolution. This technique helped identify the worm sub-groups, called clusters, that shared common phenotypes and quantified the differences between individual worms and the sub-groups. The built-in algorithm in MATLAB was used to perform the clustering analysis. The readout parameters mentioned above were standardized using Eq. 4 with their respective averages from the control worms. Each data point was assigned to a cluster by calculating the minimum Euclidean distance between the data point and the cluster centroid.

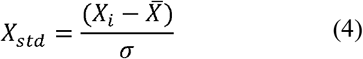

where *X_std_* is the standardized value of the data point, *X_i_* is the data point of interest (i.e., egg-count, length, diameter, length reduction, and fluorescent intensity), 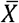 is the control sample mean, and σ is the control sample standard deviation.

Principal Component Analysis (PCA) was performed to reduce the dimensionality and obtain the parameters with the highest level of information in our datasets. Briefly, in PCA, a new set of variables called the principal components are derived by using linear combinations of the original parameters. The principal components in our studies were calculated using the Minitab software and ranked based on their decreasing eigen-values. Eigen-value represents the amount of variance in the principal component. Therefore, the first two to three principal components can explain 80% of the total variance, making them suitable for representing the entire dataset, hence reducing the dimensionality. Contributions of each original parameter towards the top principal components were calculated by normalizing the coefficients of the principal components to their L^1^ norms. The L^1^ norm is the summation of all absolute values of the coefficient of each parameter in the principal component. Using this technique, we isolated and reported the original parameters with the highest level of information.

## RESULTS AND DISCUSSIONS

In this paper, we first investigated three different designs of the microfluidic device (Figure 1) with various tapering channel sizes and quantified the worm loading and orientation efficiencies. Using the best design, we tested a new bidirectional EF stimulation protocol to ensure that the egg-laying results in the multi-worm device were comparable with the results from our single-channel device. As a proof of concept for toxicity analysis, worms were exposed to 100 mM glucose, and their electric egg count was quantified for the first time. Finally, we showed another novel application of our assay for investigating the toxicity effects of polystyrene microplastics and possible correlations between microplastic ingestion level and other on-chip phenotypes at a single-worm resolution.

### Loading efficiency of the microfluidic chip

The response of freely moving gravid adult worms to EF in a close-fitting microchannel was studied by us previously at a single worm throughput, and worms were shown to deposit eggs in a controlled manner.^27,28^ The effects of EF direction, strength, and exposure duration as well as the worms’ age and involvement of neurons and muscles in electric egg-laying were demonstrated, highlighting the potential of this method for toxicological studies at the cell to behaviour level. Here, we improved our technology by performing electric egg-laying and on-chip imaging on 8 worms in parallel, which was restricted by our microscope field of view of 2.2×1.7 mm^2^. We were also able to keep the worms’ identity known during the experiments for correlating the microplastic accumulation levels with the electric egg-laying response and some physiological parameters at a single animal resolution.

Our microfluidic device in Figure 1 was designed to (1) distribute 8 worms across the electrical traps, (2) restrain them within traps during EF stimulation, and (3) allow egg release and fluorescent imaging of the worms (Supplementary Video S1). The loading performance of our microfluidic device strongly depended on the tapered channels connecting the inlet channel network to the electrical traps (Figure 1B). They helped impeding the worms from slipping into the electric traps uncontrollably. Three microfluidic devices with tapered loading channels narrowing from a width of 60 μm into 30, 35, or 40 μm were tested for loading efficiency (Figure S2).

A total of 80 worms (in 10 trials) were tested in each device, and the tapered channel with a 30 μm wide outlet aided in smooth worm loading into the electrical traps with the highest loading efficiency of 91.25% (73/80 worms). The 35 and 40 μm-wide tapered channels showed significantly lower loading efficiencies of 56.25% (45/80 worms) and 47.5% (38/80 worms), respectively. They were large for maintaining the worms during loading while sometimes accepting more than one worm per channel. The same loading technique was used previously by Banse et al.^36^ with a similar tapered loading channel dimension of 28 μm. Thus, the rest of the experiments were conducted using the 30 μm-wide tapered channel device.

### Assay time and EF pulsation effects

In our single-worm electric egg-laying experiments^27,28^, we observed a significantly higher egg-count for worms when the anode was positioned at their anterior sides (i.e., anodal exposure). We call this method a unidirectional pulsation since we could choose the anode position with respect to the single worm orientation in this device. In a new set of experiments with our single-worm device in Figure 3A, 20 unidirectional anodal pulses (6 V/cm, 5s on, 25s off) were applied within a 10-minute experimental duration to study the egg-laying of N=32 wildtype worms. The number of eggs laid by a worm per minute dropped rapidly in the first 5 minutes and plateaued at almost no eggs per worm from minute 5 to minute 10. Figure 3B shows the total number of eggs laid per worm after 5 or 10 minutes in the single-worm device, with no statistically significant difference between the two groups (p-value>0.05). Within the first 5 minutes, the worms were exposed to 10 unidirectional anodal pulses which were selected for the rest of the experiments.

**Figure 3:**
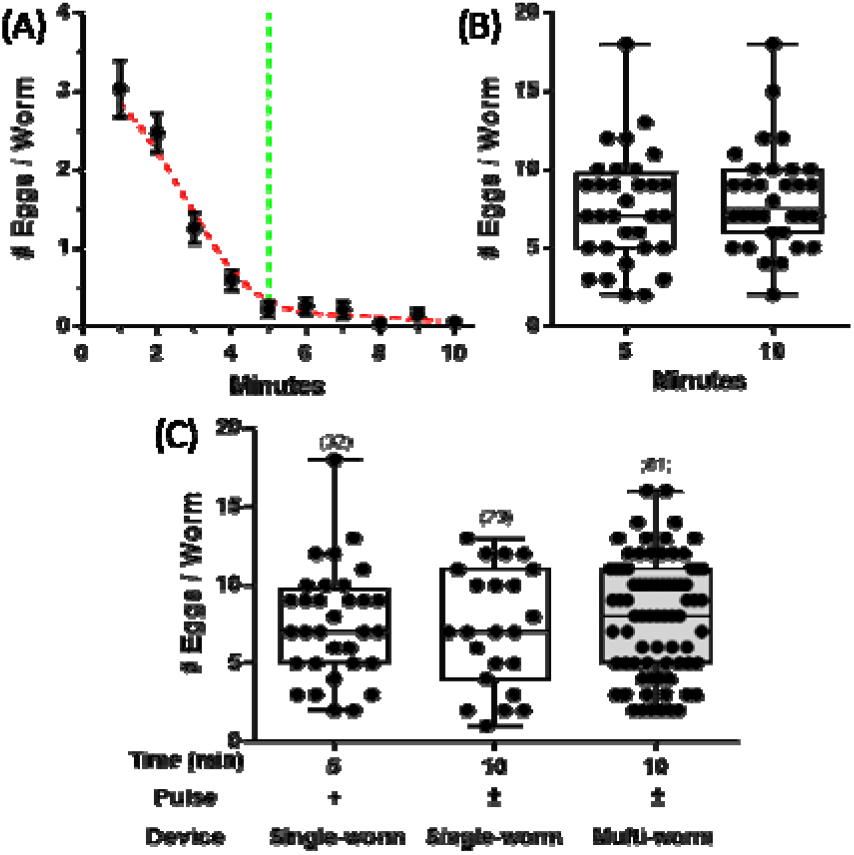
The effects of stimulation time and pulsation method on the electric egg-laying behaviour of adult C. elegans. (A) The number of eggs per worm for N=32 worms at 6V/cm using unidirectional pulsation (5s on, 25s off) inside our single-worm device^28^. (B) The total number of eggs deposited at the end of minutes 5 and 10 in (A). (C) Effects of unidirectional (+) pulsation for N=32 in the single-worm device and bidirectional (±) pulsation for N=23 and N=81 worms in the single- and multi-worm devices, respectively, studied on the number of eggs per worm during 5 and 10 minutes.

In our preliminary experiments with the multi-worm device, we noticed that the worms were oriented randomly with approximately 50% head or tail towards the trap (Figure 2 A). This prevented the use of unidirectional pulses because approximately half of the worms would have been exposed to cathodal pulses, leading to no egg-laying response and waste of animals. Similar longitudinal orientation in the multi-worm device may have been achieved by preconditioning the worms with a longitudinal stimulus (e.g., controlled EF^37^ or flow^38^), but this would have added complexity to our procedures.

We tested a new bidirectional pulsation method in order to use all the head and tail loaded worms in the multi-worm device. Accordingly, a series of 10 bidirectional EF pulses (+6V/cm for 5s, 25s off, −6V/cm for 5s, 25s off) were applied, aiming to stimulate each worm with an equal number of anodal and cathodal pulses. While this method ensured exposing each worm to 10 anodal pulses as determined in Figure 3A-3B, it also stretched the experimental duration from 5 to 10 minutes.

The effect of bidirectional pulsation with 10 anodal and 10 cathodal pulses (within 10 min) was first investigated in the single-worm device (Figure 3C, middle column). For the sake of comparison, the worms’ response to 10 unidirectional anodal pulses within 5 min in the single-worm device was also provided in Figure 3C, left column. As shown, the responses with both methods were statistically similar, proving that the cathodal exposures did not contribute significantly towards the egg-laying response in the bidirectional exposure method.

The above results ensured us that the bidirectional pulsation technique could be used in the multi-worm device to increase the throughput of the assay. The result of this experiment is also shown in Figure 3C (right grey column), depicting no statistical difference between the single-worm and multi-worm device responses while enabling us to increase the sample size from 23 to 81 worms and reduce the assay time from 4 to 2 hr, respectively.

With the multi-worm device and the bidirectional pulsation method, we were able to reach an egg-laying assay throughput of up to 40 worms/hr, which could be increased in the future with a larger microscope field of view. Compared to our electrical single-worm chip and the natural on-plate egg-laying technique, which can reach a throughput of approximately 5 worms/hr, our multi-worm assay can provide opportunities for testing more chemicals in a faster way. Next, we show the novel applications of our technique for glucose and microplastics toxicity testing.

### Effect of glucose on the electric egg-laying of *C. elegans*

*C. elegans* egg-laying circuit consists of a simple neuronal system that is serotonin-controlled. It has been used as an effective readout for identifying the effects of drugs and neurotransmitters. The egg-laying rate is affected by the culture conditions and the availability of food. For instance, different studies have shown the adverse effects of glucose on the natural egg-laying rate and life span of *C. elegans* as an application for antidiabetic drug screening.^29,39-13^ Moreover, two recent studies illustrated the associated neurotoxicity effects of glucose on protein aggregation in *C. elegans* models of Parkinson’s and Huntington’s diseases.^44,45^ Given the laboriousness and time-consuming nature of these experiments, we asked whether glucose affects the electrical egg-laying response of *C. elegans* and if this method can be used to speed up such chemical screening studies.

In each experiment, synchronized one day old adult worms were randomly picked from 100 mM glucose-dosed and un-dosed control NGM plates and loaded into the multi-worm device for testing using bidirectional pulsation at EF=6 V/cm. Their electric egg-laying count, length, diameter, and body length reduction during EF exposure were quantified, as shown in Figure 4. We also monitored the natural egg-laying behaviour of 25 worms off-chip for comparison purposes. Our egg-laying results in Figures 4A and 4B for on- and off-chip worms, respectively, showed that the control worms exhibited a strong egg-laying behaviour as illustrated by their high egg-counts in both experiments. The number of eggs/worm deposited off-chip was higher because eggs were allowed to be laid naturally over 3 hr in this experiment, which was significantly longer than the 10 min period used to electrically induce eggs on-chip. In other words, the off-chip worms had more time to reproduce new eggs, while the on-chip worms were egg-depleted rapidly and removed from the device. More importantly, the worms grown on the glucose plates showed noticeable natural and electrical egg-laying deficiencies, determined by their significantly lower egg-counts. Both on- and off-chip egg-laying experiments followed the same trends, indicating the suitability of our microfluidic technique for rapid glucose screening.

**Figure 4:**
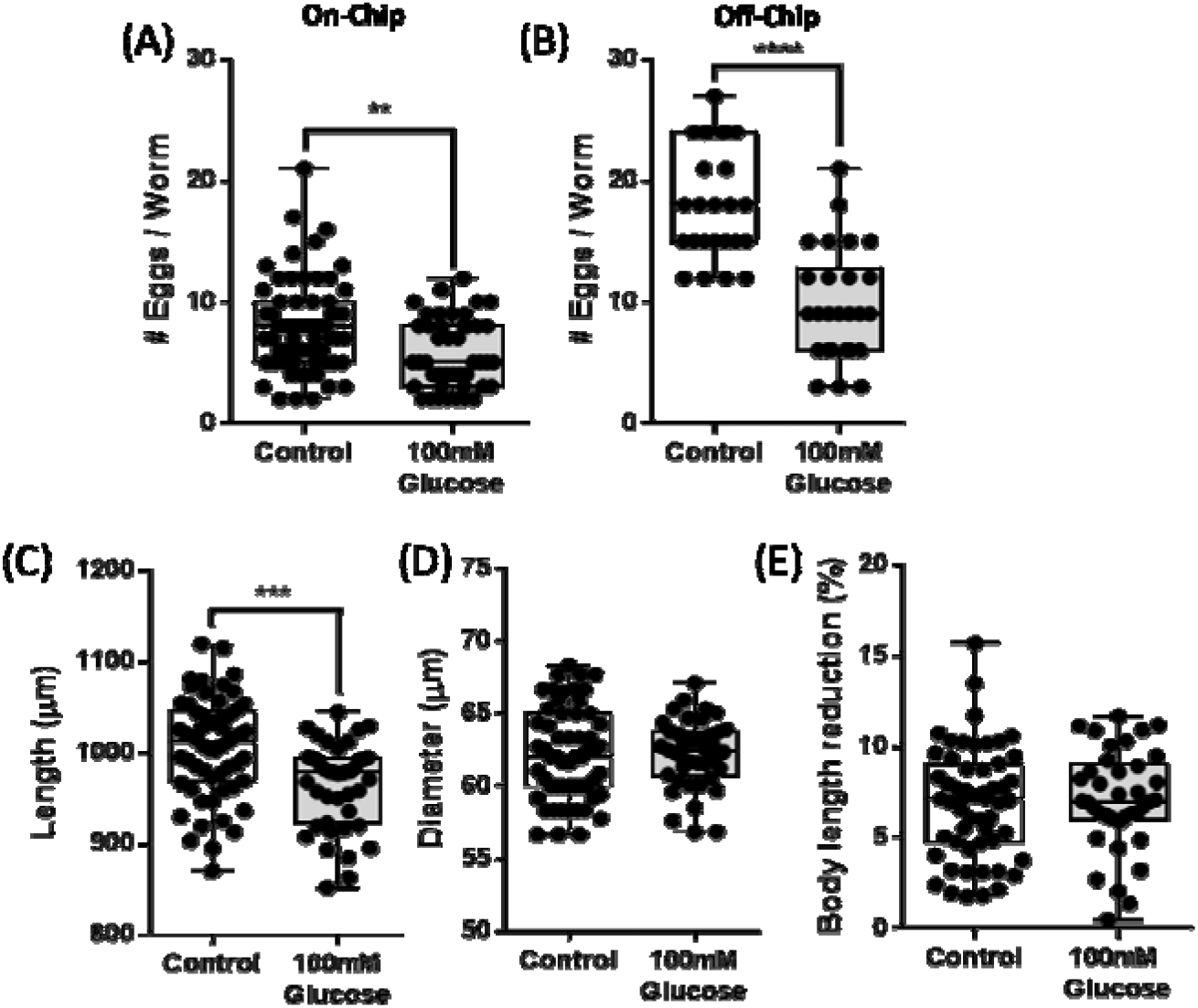
The effect of 100 mM glucose on the (A) natural off-chip (N=25) and (B) electric on-chip (N=65 and 44 for control and glucose treated worms respectively) egg-laying of adult C. elegans. EF of 6V/cm was applied for 10 min in pulses with ±5s on and 25s offperiods. Worms (C) length, (D) diameter, and (E) body length reduction during anodal stimulation were also quantified.

The off-chip experiment was not only time consuming (5 worms/hr) but also prone to the possibility of damaging the worms during the transfer process and miscounting the eggs while searching the plates. Conversely, we performed the on-chip experiments on 90-120 worms in less than three hours (30-40 worms/hr throughput) using our multi-worm device with the ability to wash the worms off the plate and into the device without having to pick and potentially damage them. Egg-laying was done on animals spatially restricted in one place, which made the assay less prone to errors in egg counting. Moreover, our technique provided not only the egg-count but also other quantitative readouts, as shown in Figure 4C-4E.

It has been shown that glucose affects the worm growth with a discrepancy in the literature, i.e., papers showing an increase^43,45^ or decrease^41,46^ in the worms’ size. Here, we investigated whether exposure to 100 mM glucose results in changes in the worms’ growth by measuring their lengths and diameters on the multi-worm device after the egg-laying assay. As shown in Figure 4C-4D, the glucose-fed worms were significantly shorter (20%) with no change in their diameter compared to the control worms. Therefore, the electric egg-laying deficiency in Figure 4B may be attributed to retardation in the growth rate.

The decrease in worms’ length might also be accompanied by possible changes in the worms’ muscle integrity. In our single-worm egg-laying experiments^28^, we showed that, during EF stimulations, worms contract their bodies in a consistent manner to release eggs. Body shortening was quantified in Figure 4E during the anodal pulses in the multi-worm device for the control and 100mM glucose-fed worms. No significant change was observed in the body length reduction during EF exposure, indicating no detectable effect of glucose on the body wall muscle activities of *C. elegans.* Yet, a deeper analysis of the contraction-relaxation mechanisms might reveal subtle phenotypes that need to be investigated in the future.

Altogether, using glucose, we provided a proof of concept for the use of our technique in determining the toxic effects of chemicals. We also showed that glucose induces abnormal electric egg-laying behaviour in *C. elegans*, which was on par with natural egg-laying behaviour reported earlier by Teshiba et al.^29^. In the future, we aim to use the technique for antidiabetic drug screening and other relevant diseases.

### Effect of microplastics on the electric egg-laying of *C. elegans*

Recently, exposure to microplastics has been shown to cause significant changes in nematodes locomotory behaviours, life span, and growth rate.^23^ Microfluidics has not been used to study the microplastics toxicity on *C. elegans*. Our device not only provides the possibility to obtain various phenotypic behaviours but also allows imaging the worms individually and keeping their identity known during an experiment. In this study, we exposed N2 worms to 100 and 1000 mg/L concentrations of 1μm red fluorescent polystyrene microparticles and investigated the accumulation of microplastics in the worms using fluorescent microscopy. The subsequent effects of microplastics on the worms’ electric egg-laying, length, diameter, and body contraction during EF exposure were also investigated.

Figure 5A shows that increasing the microplastics concentration increased the red fluorescent intensity expression in N2 *C. elegans* with microplastics accumulation in the intestine and the pharynx. The quantitative data in Figure 5B, normalized based on the signal at 1000 mg/L, demonstrates that the microplastic uptake at 100 mg/L was not significant, compared to the control worms, but increased drastically at 1000 mg/L microplastics.

**Figure 5:**
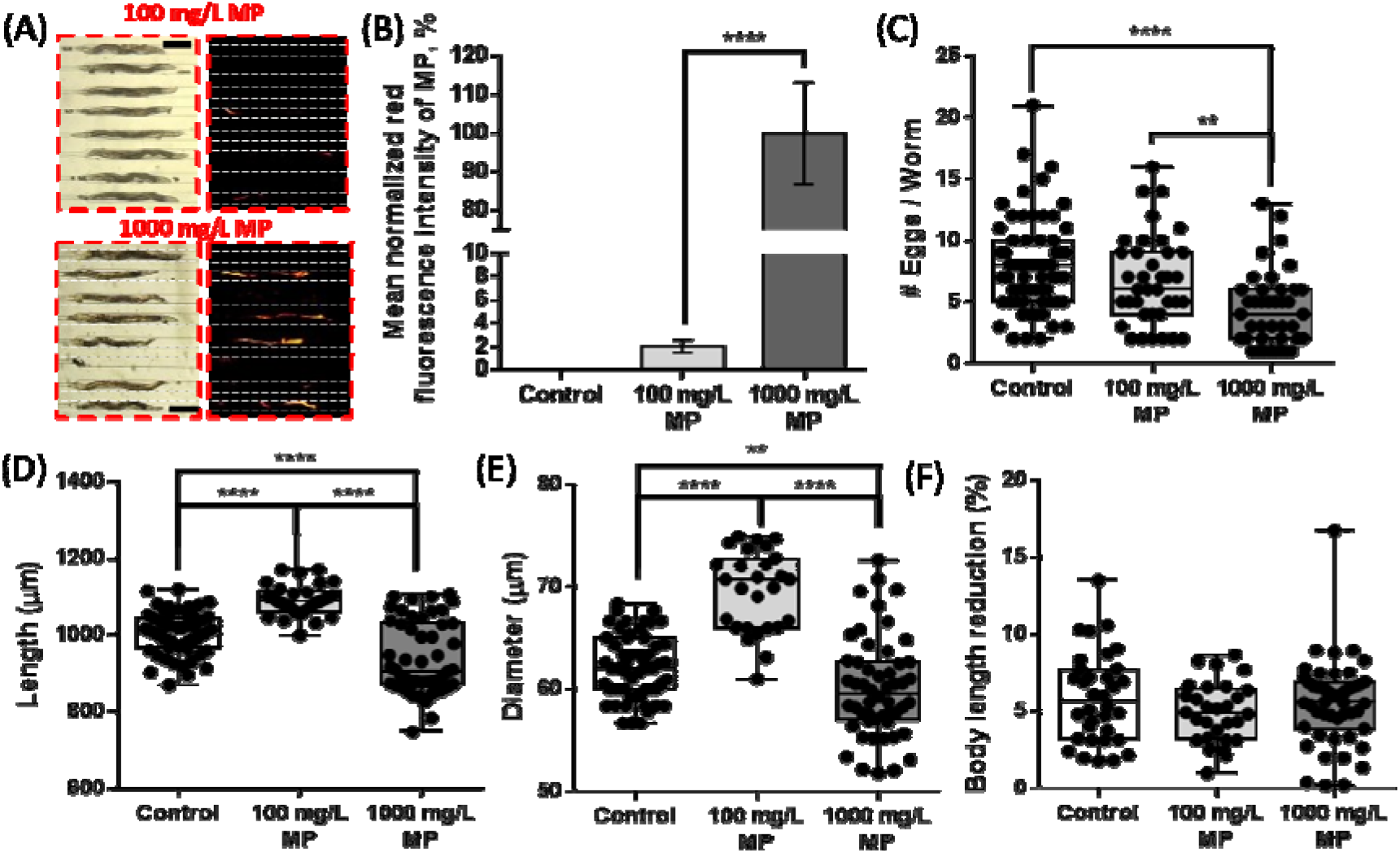
The effect of microplastics (MP) at 100 and 1000 mg/L concentrations on adult N2 C. elegans. EF=6V/cm was applied for 10 min in pulses with ±5s on and 25s off cycles. (A) Bright field and fluorescent images for the worms exposed to 100 (N= 37/39) and 1000 mg/L (N= 40/51) microplastics, indicating red fluorescent microplastic uptake (Scale bar = 200 μm). (B) Microplastic intake rate determined by calculating the mean red fluorescent intensity of the ingested microplastics and normalizing with the signal at 1000 mg/L. (C) Number of electrically deposited eggs per worm counted over 10 min. (D) Length and (E) diameter of the worms at each microplastic concentration compared to control worms (N=44). (F) Maximum total body length reduction during anodal stimulation (electrical egg-laying) in the device.

Figure 5C shows that as the microplastics concentration in the culture plate increases, the number of eggs laid electrically by a worm in our device decreases. For instance, at 100 mg/L, the microplastics uptake was relatively low (Figure 5B), resulting in a low toxic effect on egg-laying, confirmed with a slightly lower but statistically insignificant egg deposition of these worms compared to the controls, and a low number of non-responding worms (N=2/39). However, worms exposed to 1000 mg/L microplastics ingested more microplastics (Figure 5B), and their electric egg-laying was significantly lower than controls (P<0.0001, Figure 5C). It is also worth emphasizing that the number of non-responders (N = 11/51) to EF in the 1000 mg/L sample was significant which confirms the toxicity of microplastics on egg-laying.

In terms of body size, Figure 5D-5E show that the worms exposed to 1000 mg/L microplastics were significantly smaller than the control worms in length (P<0.0001) and diameter (P<0.01). However, the worms exposed to 100 mg/L microplastics were larger than controls in both length and diameter (P<0.0001). The reason behind this observation is still unknown to us and needs further investigation. Although some researchers have reported a negative effect of microplastics on *C. elegans* length^10,11,22^, others have shown similar results in terms of the non-correlating effect of microplastics on the worms’ length.^24^

Lastly, Figure 5F shows the length reduction of worms under various microplastic exposure conditions during anodal stimulations (electrical egg-laying) in our device. Length reduction did not change significantly for the treated worms compared to controls (P>0.5), indicating that microplastics did not affect the muscle contractions. We concluded that the lower egg depositions at higher microplastic concentrations, observed in Figure 5C, were potentially attributed to egg production deficiency and not the muscle activities.

### Effect of microplastics on the neuronal system of *C. elegans*

Two recent studies^11,47^ have reported neuronal damages associated with nano- and micro-plastics exposure in *C. elegans* GABAergic, cholinergic, and dopaminergic neurons, suggesting that the locomotory deficiencies may be due to neurotoxicity. Our goal was to investigate the relationship between the microplastics induced neurotoxicity, growth deficiency, and electric egg-laying using NW1229, a transgenic strain expressing GFP pan-neuronally. NW1229 worms were exposed to 100 and 1000 mg/L of 1 μm red fluorescent polystyrene particles. They were then loaded into the device, and their microplastics intake (red fluorescence), electric egg count, length, diameter, length reduction during anodal stimulation, and GFP expression of the neurons were investigated. The results are shown in Figures 6 and 7.

**Figure 6:**
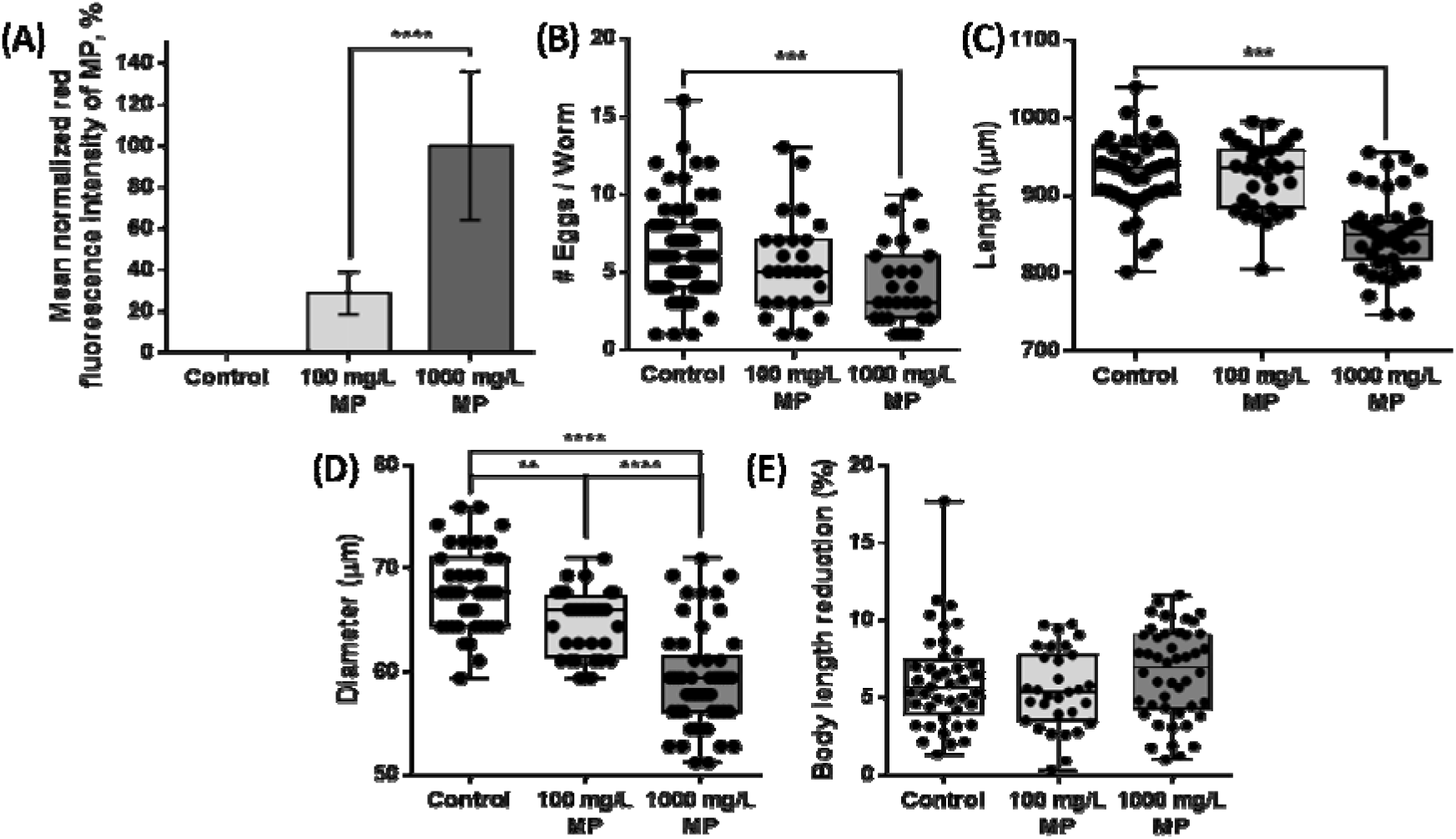
The effect of microplastics (MP) at 100 mg/L (N =26/30) and 1000 mg/L (N= 27/45) concentrations on adult NW1229 C. elegans expressing GFP pan-neuronally. EF=6V/cm was applied for 10 min in pulses with ±5s on and 25s off cycles. (A) Microplastic intake rate determined by calculating the mean red fluorescent intensity of the ingested microplastics and normalizing with the signal at 1000 mg/L. (B) Number of electrically deposited eggs per worm counted over 10 min. (C) Length and (D) diameter of the worms at each microplastic concentration compared to control worms (N = 41). (E) Maximum total body length reduction during anodal stimulation (electric egg-laying) in the device.

Results in Figure 6A showed similar trends to the microplastics uptake by N2 worms (Figure 5B). Control worms not exposed to microplastics did not express any red fluorescent while microplastics ingested at 1000 mg/L concentration was significantly higher than 100 mg/L (P<0.0001). Our egg-laying results in Figure 6B showed that the untreated worms exhibited a strong electric egg-deposition, as demonstrated by their high egg count. However, the worms treated with microplastics exhibited noticeable egg-laying deficiency at 1000 mg/L concentration (P<0.001). At 100 mg/L, microplastics had a slight effect on egg deposition, which was not significantly different from controls, just like what we observed for N2 worms. Also, the number of non-responders (N = 18/45) to EF in the 1000 mg/L sample was higher than the 100 mg/L (N =4/30). In terms of body size, Figures 6C-6D show that the worms exposed to 100 mg/L microplastics were similar in length (P> 0.5) but significantly thinner in diameter (P<0.01) compared to the control worms. This was contrary to what we observed for the N2 strain, which requires further investigation in the future. The worms treated with 1000 mg/L microplastics were significantly shorter (P<0.0001) and thinner (P<0.0001) than the control worms and the 100 mg/L treated worms. Finally, we confirmed that the anodal length reductions for all conditions were consistent and similar to our observation for the N2 worms, indicating that the length reduction was not significantly (P>0.2) altered due to exposure to microplastics (Figure 6E).

**Figure 7:**
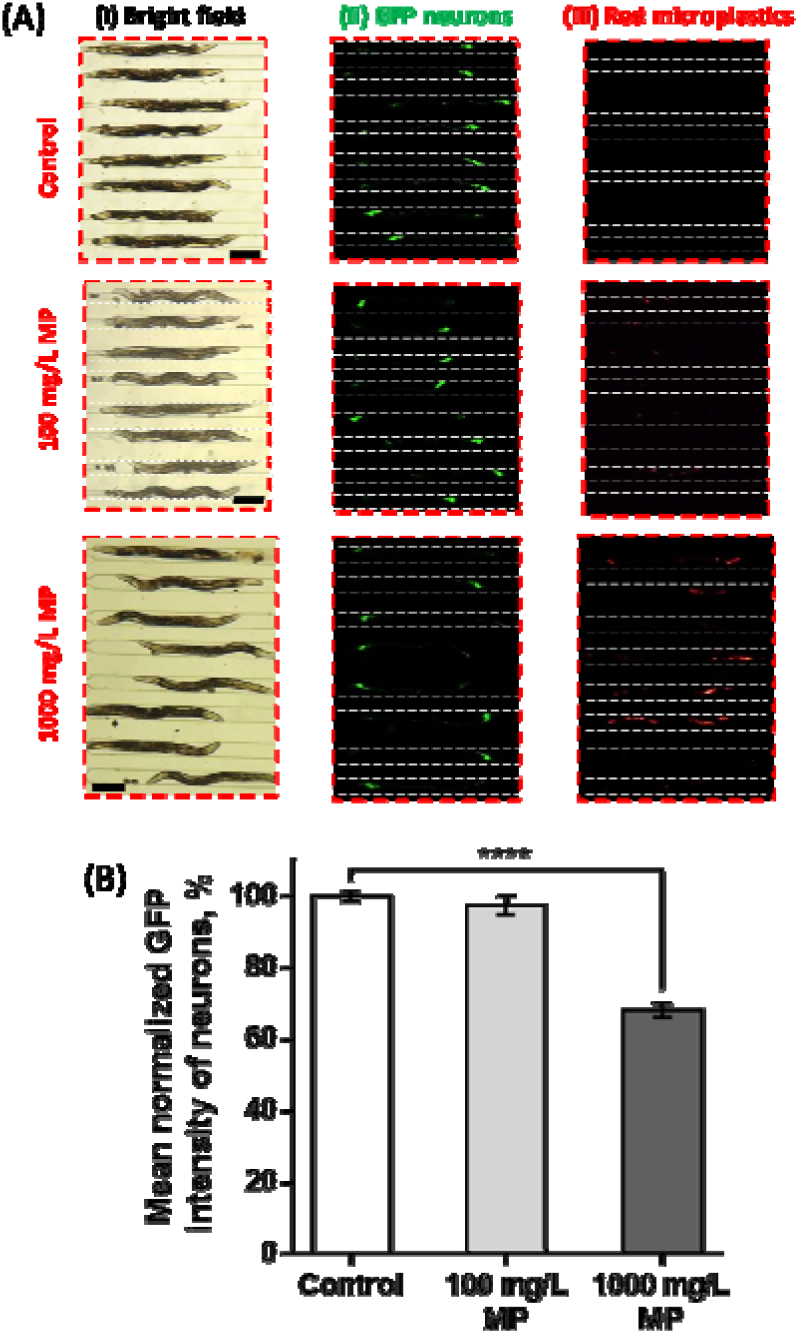
GFP expression of NW1229 C. elegans fed with 100 and 1000 mg/L. (A) Normalized mean fluorescent intensities. (B) Bright field (Scale bar = 200 μm) (i) and fluorescent images for neuronal system (ii) and microplastics (iii) at 0 (control), 100, and 1000 mg/L microplastics.

To investigate neurotoxicity, following the 10 min egg-laying experiment, GFP expressing neurons and red fluorescent microparticles in the worms were separately imaged in the electric traps (Figure 7A). The mean GFP intensities of neurons at both microplastic concentrations were normalized by the average GFP value obtained for the control worms grown without microplastics (Figure 7B). Similar to the electric egg-laying results, exposure to 100 mg/L microplastics resulted in insignificant (P>0.6) GFP decay compared to the control worms. However, a significant decrease (P <0.0001) of approximately 35% in GFP expression was obtained at 1000 mg/L microplastic exposure (Figure 7B), similar to the egg-count reduction in the same condition (Figure 6B). The decreasing trend in GFP expression at both concentrations is in line with the increasing trend in microplastics uptake, shown in Figure 6A for the NW1229 worms. The above results show the appropriateness of our technique for toxicity assessment from neuron to behaviour level. Moreover, our results confirmed that microplastics toxicity is not specific towards certain neurons, whereas GFP expression decayed over the entire body of the worm. This was also confirmed previously using off-chip assays that showed neurodegeneration for specific neurons as well as defective locomotory behaviour.^23^

Altogether, our results showed the suitability of our microfluidic technique to study glucose and microplastics toxicity in *C. elegans* and detect the associated behavioural and neuronal abnormalities, quantitively and at a population resolution. The following section will show the potential of our device for use in single-worm level neurobehavioural toxicity studies.

### Single-worm analysis reveals heterogeneity in microplastics uptake and correlating egg-laying and physiological phenotypes

In recent years, physicians have found that drug-patient interactions are highly important, demonstrating the need for developing individualized diagnoses and treatments that take patient variabilities into account^48^. In the case of *C. elegans*, multiple microfluidic devices have recently shown that analysing the data in an individual-based manner would reveal subtle phenotypes that were hidden in population measurements^49,50^. Here, we questioned the heterogeneity of microplastics uptake and toxicity, and show that our microfluidic device can be used to perform phenotypical and neuronal analyses at single-animal resolution and at a throughput higher than conventional assays for microplastics toxicity assessment.

As described in the Materials and Methods section, HCA and PCA were performed on individual N2 and NW1229 worms treated with microplastics at 100 and 1000 mg/L, and the results were compared with their counterpart control worms. The egg-count, worm length, worm diameter, length reduction during anodal stimulation, and normalized mean GFP and RFP fluorescent intensities, representing neurons and microplastics, respectively, were quantified for individual worms.

Figure 8Ai shows the heatmap representing the clustering generated by the HCA algorithm for N2 worms. Columns and rows represent individual worms and their phenotypes, respectively. The data were standardized using Eq.4 based on the control worm parameters. Thus, in Figure 8Ai, the color gradients depict whether a parameter of interest is quantitatively similar (white), lower (dark blue), or higher (dark red) than the average of the control group. The only exemption was the microplastic uptake RFP parameter for which the 100 mg/L data was selected as standardization reference since the control worms were not exposed to microplastics, expressing zero RFP intensity.

**Figure 8:**
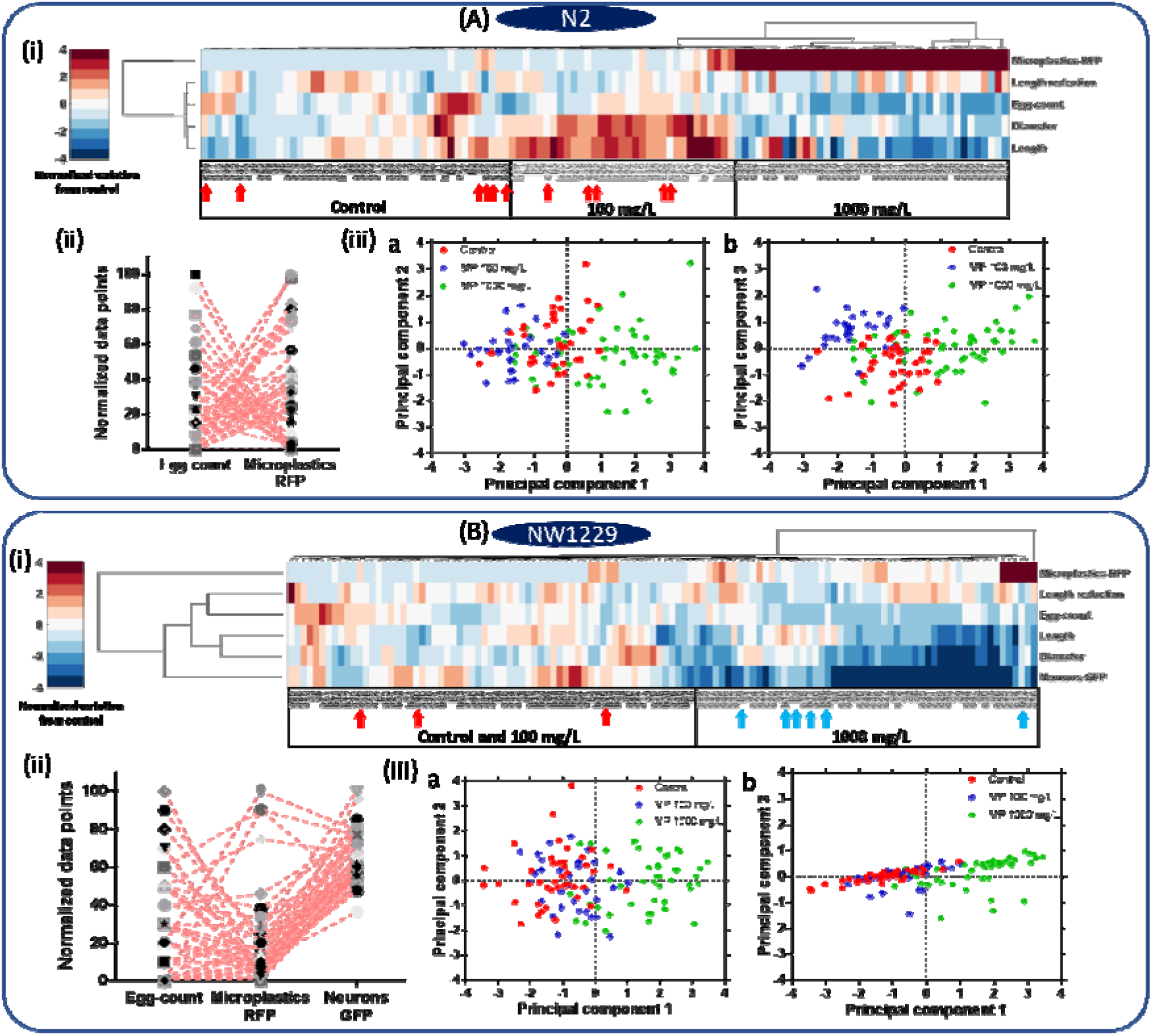
Phenotypic analysis of individual worms, (A) N2 and (B) NW1229, including control worms and ones treated with 100 and 1000 mg/L microplastics. (i) Hierarchical Cluster Analysis (HCA) of different phenotypes of the control and microplastic-exposed worms. (ii) Normalized individual responses of egg-count, microplastic uptake and GFP expression (NW1229 strain only) at 1000 mg/L. (iii) Principal Component Analysis (PCA) of control, 100 and 1000 mg/L microplastic-exposed worms.

Figure 8Ai shows that the control worms were mostly clustered at the left-hand side of the graph, followed by the worms exposed to 100 and 1000 mg/L microplastics. Generally, the graph confirms the population trends discussed earlier in the paper, but at an individual worm level. It shows that the microplastics uptake at 1000 mg/L was much higher than the 100 mg/L, for most of the worms, and the worms’ diameter and length at 1000 mg/L were lower than the control and 100 mg/L worms. Moreover, the electric egg-laying rate for the worms that took more microplastics was always lowered. Interestingly, using the clustering technique, we were able to isolate individual worms treated with 1000 mg/L microplastics that were behaving like untreated worms, with high egg count and low microplastics accumulation. These worms were clustered within the control or 100 mg/L populations as depicted with red arrows in Figure 8Ai. This showed the heterogeneity of microplastic uptake in the worms treated in the same way, which necessitates single-animal resolution assays like the one offered here to isolate such worms and understand their correlating phenotypes correctly. Such data points were hidden in our population-based averaging results offered in the previous sections, and this is the case in conventional worm screening assays, especially when they involve behavioral phenotyping.

HCA and PCA revealed some hidden correlations between the independent parameters. For instance, using PCA, the contribution of each parameter to the first, second, and third principal components, computed in Table S1, showed that the egg-count and the microplastics uptake level are inversely proportional. In other words, worms with higher microplastics expression were defective in egg-laying. To better visualize this correlation, we focused on the 1000 mg/L treated worms and normalized the egg-count and microplastics fluorescent expression parameters to obtain similar scales for comparison purposes. Then we plotted both parameters while preserving the identity of individual worms by connecting dash-lines, as shown in Figure 8Aii. Moreover, our results were further confirmed by plotting the calculated scores from the PCA using the first, second, and third principal components (Figure 8Aiii). This figure also confirms that the 1000 mg/L treated worms were clustered together with some individuals behaving like control and existing within the control cluster as discussed above.

Similar results were obtained for the NW1229 strain as shown in Figure 8B. Briefly, HCA analysis in Figure 8Bi created two main clusters for the 1000 mg/L worms at the right-hand side and the control and 100 mg/L worms mixed at the left-hand side, explaining their population trends described earlier. The heatmap illustrates increase in the microplastic uptake at 1000 mg/L exposure, and corresponding decreases in the egg-count, length, diameter, and neurons GFP expression of the worms. The graph also shows that there was a large variation in microplastics uptake within the 1000 mg/L population with some worms showing a significantly higher uptake than the others. Moreover, some NW1229 worms, exposed to 100 mg/L (blue arrows) and 1000 mg/L (red arrows) microplastics, were not placed in their respective clusters in Figure 8Bi. For these worms, the level of microplastics uptake (or GFP expression) was on par with the same phenotypes in their clusters.

The PCA analysis in Table S2 for NW1229 worms showed similar trends to the N2 strain, demonstrating that the microplastics uptake and egg-count were inversely proportional. However, there was no obvious pattern between the microplastics uptake and neurodegeneration. This was observed in Figure 8Bii by plotting the individual responses of 1000 mg/L worms for the egg-count, microplastics uptake, and neurons fluorescent expression while preserving the worms’ identity. Lastly, Figure 8Biii shows that only using the first and second principal components, our results were further confirmed, and the 1000 mg/L treated worms were clustered together, whereas the control and 100 mg/L treated worms were mixed.

All in all, we envision that our microfluidic device, coupled with the clustering techniques above, would be of benefit to develop individualized drug and pollutant screening assays for *C. elegans*.

## Conclusion

As an emerging toxicant, microplastics have been recently spotted in abundance throughout the environment. *C. elegans* is a simple and easy to use model organism for toxicological studies. Previous studies showed the negative effects of microplastics on *C. elegans* phenotypical behaviours using conventional assays, which are time-consuming and laborious, limiting the possibility of developing high throughput toxicity screening methods. Moreover, previous studies have solely focused on the natural behaviours of *C. elegans* while paying less attention to the sensory-motor evoked phenotypes to assess the neuronal system integrity. Therefore, in this paper, we developed a microfluidic device to exploit the newly introduced *C. elegans* electric egg-laying behaviour for toxicity studies.

Our device enabled trapping and exposing up to 8 individual worms in parallel to EF while allowing on-chip fluorescent imaging. We optimized our microfluidic device and adopted a new bidirectional stimulation technique to achieve a throughput of 40 worms/hr, which can be further enhanced in the future with a larger microscope field of view. Large datasets of multiple parameters such as electric egg-count, worm length, worm diameter, length reduction, and neuronal and microplastics mean fluorescent intensities were obtained and analyzed in population-averaged and individual-worm approaches.

The effect of glucose was investigated as a proof of concept for toxicity assays. Our findings implied that 100 mM of glucose induced abnormal electric egg-laying behaviour and resulted in a smaller body size. Thus, in the future, we aim to use the technique for antidiabetic drug screening. Moreover, we employed our device with two *C. elegans* strains (N2 and NW1229) to investigate the toxicity effects of 1 μm polystyrene microparticles at concentrations of 100 and 1000 mg/L on electric egg-laying and worm size as well as neurodegeneration in a faster manner using single-worm phenotyping. Our results showed that exposing the worms to 1000 mg/L caused severe egg-laying deficiency and neurodegeneration, and reduction in body size. Lastly, using HCA and PCA, we showed that single-worm phenotyping could reveal heterogeneity in microplastics uptake, which was correlated with the deficiency in egg-laying.

In the future, we aim to develop a microfluidic device for life-long phenotyping of microplastics toxicity while allowing for sensory-motor behavioural studies. Our technique can also be used as an ecotoxicity screening technique for determining the sublethal effects of other materials such as heavy metals on *C. elegans*.

## Author Contributions

**Khaled Youssef**: Methodology, Investigation, Formal analysis, Validation, Data curation, Visualization, Writing - original draft. **Daphne Archonta**: Data curation and analysis. **Terrance J. Kubiseski**: Conceptualization, Writing - review & editing. **Anurag Tandon**: Supervision, Validation, Writing - review & editing. **Pouya Rezai**: Conceptualization, Methodology, Validation, Resources, Writing - review & editing, Supervision, Funding acquisition.

## Acknowledgements

This work was supported by Natural Sciences and Engineering Research Council (NSERC) of Canada and the Early Researcher Award to PR and the Ontario Trillium Scholarship to KY.

## References

1. Wright SL, Kelly FJ. Plastic and Human Health: A Micro Issue? Environmental Science and Technology. 2017;51(12):6634–6647. doi:10.1021/acs.est.7b00423

2. Geyer R, Jambeck JR, Law KL. Production, use, and fate of all plastics ever made. Science Advances. 2017;3(7):e1700782. doi:10.1126/sciadv.1700782

3. Sharma S, Chatterjee S. Microplastic pollution, a threat to marine ecosystem and human health: a short review. Environmental Science and Pollution Research. 2017;24(27):21530–21547. doi:10.1007/s11356-017-9910-8

4. Sussarellu R, Suquet M, Thomas Y, et al. Oyster reproduction is affected by exposure to polystyrene microplastics. Proceedings of the National Academy of Sciences. 2016;113(9):2430 LP–2435. doi:10.1073/pnas.1519019113

5. Prüst M, Meijer J, Westerink RHS. The plastic brain: neurotoxicity of micro- and nanoplastics. Particle and Fibre Toxicology. 2020;17(1):24. doi:10.1186/s12989-020-00358-y

6. PlasticsEurope (PEMRG). (September 25, 2019). Production of plastics worldwide from 1950 to 2018 (in million metric tons)* [Graph]. In Statista. Retrieved December 09, 2020, from https://www.statista.com/statistics/282732/global-production-of-plastics-sin.

7. de Sá LC, Oliveira M, Ribeiro F, Rocha TL, Futter MN. Studies of the effects of microplastics on aquatic organisms: What do we know and where should we focus our efforts in the future? Science of the Total Environment. 2018;645:1029–1039. doi:10.1016/j.scitotenv.2018.07.207

8. Pennino MG, Bachiller E, Lloret-Lloret E, et al. Ingestion of microplastics and occurrence of parasite association in Mediterranean anchovy and sardine. Marine Pollution Bulletin. 2020;158:111399. doi:10.1016/j.marpolbul.2020.111399

9. Kim SW, Kim D, Jeong SW, An YJ. Size-dependent effects of polystyrene plastic particles on the nematode Caenorhabditis elegans as related to soil physicochemical properties. Environmental Pollution. 2020;258:113740. doi:10.1016/j.envpol.2019.113740

10. Shang X, Lu J, Feng C, et al. Microplastic (1 and 5 μm) exposure disturbs lifespan and intestine function in the nematode Caenorhabditis elegans. Science of the Total Environment. 2020;705:135837. doi:10.1016/j.scitotenv.2019.135837

11. Lei L, Liu M, Song Y, et al. Polystyrene (nano)microplastics cause size-dependent neurotoxicity, oxidative damage and other adverse effects in Caenorhabditis elegans. Environmental Science: Nano. 2018;5(8):2009–2020. doi:10.1039/c8en00412a

12. Lei L, Wu S, Lu S, et al. Microplastic particles cause intestinal damage and other adverse effects in zebrafish Danio rerio and nematode Caenorhabditis elegans. Science of the Total Environment. 2018;619-620:1–8. doi:10.1016/j.scitotenv.2017.11.103

13. Olivier K, Karanth S. Toxicology testing: in vivo mammalian models. In: An Introduction to Interdisciplinary Toxicology. Elsevier; 2020:487–506. doi:10.1016/b978-0-12-813602-7.00035-1

14. Yong CQY, Valiyaveetill S, Tang BL. Toxicity of Microplastics and Nanoplastics in Mammalian Systems. International Journal of Environmental Research and Public Health. 2020;17(5):1509. doi:10.3390/ijerph17051509

15. Hunt PR. The C. elegans model in toxicity testing. Journal of Applied Toxicology. 2017;37(1):50–59. doi:10.1002/jat.3357

16. Rand MD, Montgomery SL, Prince L, Vorojeikina D. Developmental Toxicity Assays Using the *Drosophila* Model. Current Protocols in Toxicology. 2014;59(1):1.12.1–1.12.20. doi:10.1002/0471140856.tx0112s59

17. Rubinstein AL. Zebrafish assays for drug toxicity screening. Expert Opinion on Drug Metabolism and Toxicology. 2006;2(2):231–240. doi:10.1517/17425255.2.2.231

18. Caballero MV, Candiracci M. Zebrafish as screening model for detecting toxicity and drugs efficacy. Journal of Unexplored Medical Data. 2018;3(2):4. doi:10.20517/2572-8180.2017.15

19. Youssef K, Bayat P, Peimani AR, Dibaji S, Rezai P. Miniaturized Sensors and Actuators for Biological Studies on Small Model Organisms of Disease. In: Environmental, Chemical and Medical Sensors. Springer; 2018:199–225. doi:10.1007/978-981-10-7751-7_9

20. Youssef K, Tandon A, Rezai P. Studying Parkinson’s disease using Caenorhabditis elegans models in microfluidic devices. Integrative biology□: quantitative biosciences from nano to macro. 2019;11(5):186–207. doi:10.1093/intbio/zyz017

21. Gupta BP, Rezai P. Microfluidic approaches for manipulating, imaging, and screening C. elegans. Micromachines. 2016;7(7):123. doi:10.3390/mi7070123

22. Yu Y, Chen H, Hua X, et al. Polystyrene microplastics (PS-MPs) toxicity induced oxidative stress and intestinal injury in nematode Caenorhabditis elegans. Science of the Total Environment. 2020;726:138679. doi:10.1016/j.scitotenv.2020.138679

23. Hu J, Li X, Lei L, Cao C, Wang D, He D. The Toxicity of (Nano)Microplastics on C. elegans and Its Mechanisms. In:He D, Luo Y, eds. Microplastics in Terrestrial Environments: Emerging Contaminants and Major Challenges. Cham: Springer International Publishing; 2020:259–278. doi:10.1007/698_2020_452

24. Schöpfer L, Menzel R, Schnepf U, et al. Microplastics Effects on Reproduction and Body Length of the Soil-Dwelling Nematode Caenorhabditis elegans. Frontiers in Environmental Science. 2020;8:41. doi:10.3389/fenvs.2020.00041

25. Kim Y, Jeong J, Lee S, Choi I, Choi J. Identification of adverse outcome pathway related to high-density polyethylene microplastics exposure: Caenorhabditis elegans transcription factor RNAi screening and zebrafish study. Journal of Hazardous Materials. 2020;388:121725. doi:10.1016/j.jhazmat.2019.121725

26. Maulik M, Mitra S, Bult-Ito A, Taylor BE, Vayndorf EM. Behavioral phenotyping and pathological indicators of Parkinson’s disease in C. elegans models. Frontiers in Genetics. 2017;8(JUN):77. doi:10.3389/fgene.2017.00077

27. Youssef K, Archonta D, Kubiseski TJ, Tandon A, Rezai P. Electric Egg-Laying: Effect of Electric Field in a Microchannel on C. elegans Egg-Laying Behavior. bioRxiv. 2020.

28. Youssef K, Archonta D, Kubiseski TJ, Tandon A, Rezai P. Electric Egg-Laying: Effect of Electric Field in a Microchannel on C. elegans Egg-Laying Behavior. Lab on a Chip. 2021:Under review.

29. Teshiba E, Miyahara K, Takeya H. Glucose-induced abnormal egg-laying rate in Caenorhabditis elegans. Bioscience, Biotechnology, and Biochemistry. 2016;80(7):1436–1439. doi:10.1080/09168451.2016.1158634

30. Stiernagle T. Maintenance of C. elegans. C elegans. 1999;2:51–67.

31. Porta-de-la-Riva M, Fontrodona L, Villanueva A, Cerón J. Basic Caenorhabditis elegans methods: Synchronization and observation. Journal of Visualized Experiments. 2012;(64):e4019. doi:10.3791/4019

32. Hulme SE, Shevkoplyas SS, Apfeld J, Fontana W, Whitesides GM. A microfabricated array of clamps for immobilizing and imaging C. elegans. Lab on a Chip. 2007;7(11):1515–1523. doi:10.1039/b707861g

33. Aryasomayajula A, Bayat P, Rezai P, Selvaganapathy PR. Microfluidic Devices and Their Applications. In: Springer Handbook of Nanotechnology. Springer; 2017:487–536.

34. Xia Y, Whitesides GM. Soft lithography. Annual Review of Materials Science. 1998;28(1):153–184. doi:10.1146/annurev.matsci.28.1.153

35. Youssef K, Archonta D, Kubiseski T, Tandon A, Rezai P. On-Demand Electric Field Induced Egg Laying of CAENORHABDITIS ELEGANS. In: Mtas 2019. Basel, Switzerland; 2019:392–393.

36. Banse SA, Blue BW, Robinson KJ, Jarrett CM, Phillips PC. The Stress-Chip: A microfluidic platform for stress analysis in Caenorhabditis elegans.Kim H, ed. PLOS ONE. 2019;14(5):e0216283. doi:10.1371/journal.pone.0216283

37. Youssef K, Archonta D, Kubiseski T, Tandon A, Rezai P. Parallel-Channel Electrotaxis and Neuron Screening of Caenorhabditis elegans. Micromachines. 2020;11(8):756.

38. Ge A, Wang X, Ge M, et al. Profile analysis of: C. elegans rheotaxis behavior using a microfluidic device. Lab on a Chip. 2019;19(3):475–483. doi:10.1039/c8lc01087k

39. Schulz TJ, Zarse K, Voigt A, Urban N, Birringer M, Ristow M. Glucose Restriction Extends Caenorhabditis elegans Life Span by Inducing Mitochondrial Respiration and Increasing Oxidative Stress. Cell Metabolism. 2007;6(4):280–293. doi:10.1016/j.cmet.2007.08.011

40. Schlotterer A, Kukudov G, Bozorgmehr F, et al. C. elegans as model for the study of high glucose-mediated life span reduction. Diabetes. 2009;58(11):2450–2456. doi:10.2337/db09-0567

41. Wang X, Zhang L, Zhang L, et al. Effects of excess sugars and lipids on the growth and development of Caenorhabditis elegans. Genes and Nutrition. 2020;15(1):1. doi:10.1186/s12263-020-0659-1

42. Lee SJ, Murphy CT, Kenyon C. Glucose Shortens the Life Span of C. elegans by Downregulating DAF-16/FOXO Activity and Aquaporin Gene Expression. Cell Metabolism. 2009;10(5):379–391. doi:10.1016/j.cmet.2009.10.003

43. Alcántar-Fernández J, Navarro RE, Salazar-Martínez AM, Pérez-Andrade ME, Miranda-Ríos J. Caenorhabditis elegans respond to high-glucose diets through a network of stress-responsive transcription factors.Dupuy D, ed. PLOS ONE. 2018;13(7):e0199888. doi:10.1371/journal.pone.0199888

44. Landon G, Whitney W, Priya R, Mindy F. Glucose effects on polyglutamine-induced proteotoxic stress in Caenorhabditis elegans. Biochemical and Biophysical Research Communications. 2020;522(3):709–715. doi:10.1016/j.bbrc.2019.11.159

45. Guzman AC DE, Kim EJ, Cho JH, Kim JH, Choi SS. High Glucose Diet Attenuates Dopaminergic Neuronal Function in C. elegans Leading to Acceleration of Aging Process. May 2020. doi:10.21203/rs.3.rs-29098/v1

46. Lei W, Beaudoin-Chabot C, Thibault G. Glucose increases the lifespan of post-reproductive C. elegans independently of FOXO. bioRxiv. June 2018:347435. doi:10.1101/347435

47. Qu M, Kong Y, Yuan Y, Wang D. Neuronal damage induced by nanopolystyrene particles in nematode Caenorhabditis elegans. Environmental Science: Nano. 2019;6(8):2591–2601. doi:10.1039/C9EN00473D

48. Schork NJ. Personalized medicine: time for one-person trials. Nature. 2015;520(7549):609–611.

49. Letizia MC, Cornaglia M, Trouillon R, et al. Microfluidics-enabled phenotyping of a whole population of C. elegans worms over their embryonic and post-embryonic development at single-organism resolution. Microsystems & Nanoengineering. 2018;4(1):6. doi:10.1038/s41378-018-0003-8

50. Kopito RB, Levine E. Durable spatiotemporal surveillance of Caenorhabditis elegans response to environmental cues. Lab on a Chip. 2014;14(4):764–770. doi:10.1039/c3lc51061a

