## Supplementary file for "Microfluidic Electric Egg-Laying Assay and Application to *In-vivo* Toxicity Screening of Microplastics using *C. elegans*"

**S1. Numerical Simulation of Electric Field in the Microfluidic Device**

A two dimensional (2D) numerical simulation was conducted to obtain the electric field (EF) distribution in the microfluidic device. The steady-state direct-current (DC) module of the COMSOL Multiphysics® software was used to determine the EF using Ohm’s law. The computer-aided design (CAD) software SOLIDWORKS® was used to generate the computational domain that was imported into COMSOL for mesh generation. The microfluidic device contained 8 parallel worm-dwelling microchannels called electric traps in which the worms were electrically stimulated for egg deposition (Figure 1 of the paper). Each electric trap was 85 µm-wide and 1.3 mm long. The electric traps had tapering channels at their anterior-posterior sides which were connected to the inlet and outlet channels via tree-like branching channels. The boundary conditions included an electric insulation applied to all boundaries, and electric potentials of 34V and ground applied at the two end reservoirs. For mesh generation, the triangular automatic mesh generation module of COMSOL was utilized. The EF distribution within the microfluidic device determined through the simulation is depicted in Figure 1 of the paper.

**S2. Supplementary Figures**

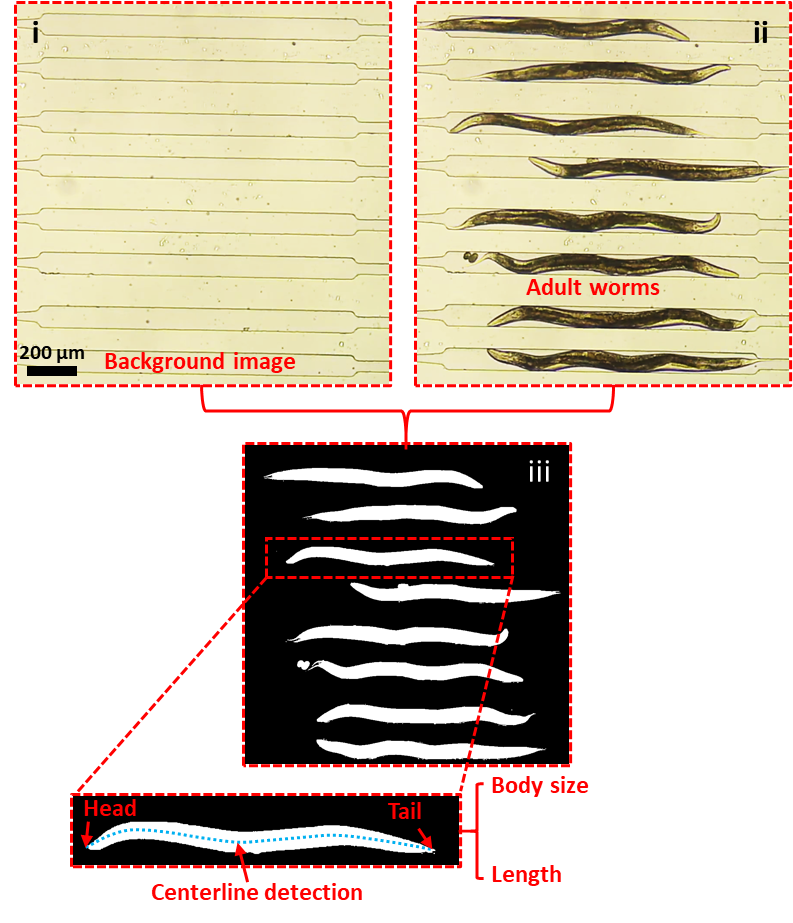

Figure S1: Data analysis by MATLAB image processing. Background subtraction was done using images (i) and (ii) to obtain the binary image (iii). (iii) Binarized image showing all worms with an inset of a single worm that was used for centerline detection and analysis of body length and length reduction.

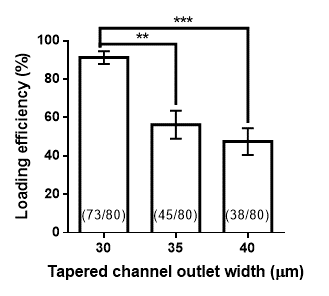

Figure S2: Worm loading efficiency of the 8 electric traps in three microfluidic devices tested with different tapered channel end-widths of 30, 35, and 40 µm. A total of 80 worms in 10 trials were tested in each device.

**S3. Supplementary Videos**

**Video S1:** Worm loading, egg-laying, and neuronal and microplastics imaging in the microfluidic device.
